# *Drosophila* CLAMP is an essential protein with sex-specific roles in males and females

**DOI:** 10.1101/042820

**Authors:** Jennifer A. Urban, Caroline A. Doherty, William T. Jordan, Jacob E. Bliss, Jessica Feng, Marcela M. Soruco, Leila E. Rieder, Erica N. Larschan

## Abstract

Dosage compensation is a fundamental mechanism in many species that corrects for the inherent imbalance in X-chromosome copy number between XY males and XX females. In *Drosophila melanogaster*, transcriptional output from the single male X-chromosome is equalized to that of XX females by recruitment of the Male Specific Lethal (MSL) complex to specific sequences along the length of the X-chromosome. The initial recruitment of MSL complex to the X-chromosome is dependent on a recently discovered zinc finger protein called Chromatin-Linked Adapter for MSL Proteins (CLAMP). However, further studies on the *in vivo* function of CLAMP remained difficult because the location of the gene in pericentric heterochromatin made it challenging to create null mutations or deficiencies. Using the CRISPR/Cas9 genome editing system, we generated the first null mutant in the *clamp* gene that eliminates expression of CLAMP protein. We show that CLAMP is necessary for both male and female viability. While females die at the third instar larval stage, males die earlier, likely due to the essential role of CLAMP in male dosage compensation. Moreover, we demonstrate that CLAMP promotes dosage compensation in males and represses key male-specific transcripts involved in sex-determination in females. Our results reveal that CLAMP is an essential protein with dual roles in males and females, which together assure that dosage compensation is a sex-specific process.

## Introduction

Many species employ a sex determination system that generates an inherent imbalance in sex chromosome copy number, such as the XX/XY system in most mammals and some insects. In this system, one sex within the species has twice the number of X-chromosome encoded genes compared to the other. Therefore, a mechanism of dosage compensation is required to equalize levels of X-linked transcripts, both between sexes and between the X and autosomes (Lucchesi et al., 2005). Dosage compensation is an essential mechanism that corrects for this imbalance by coordinately regulating gene expression of most X-chromosome encoded genes.

In *Drosophila, melanogaster*, transcription from the single male X-chromosome is increased twofold by recruitment of the Male Specific Lethal (MSL) complex. MSL complex is composed of two structural proteins, MSL1 and MSL2, three accessory proteins, MSL3, MOF (Males absent On the First), and MLE (Maleless), as well as two structural and functionally redundant non-coding RNAs, *roX1* (RNA on the X) and *roX2* (Meller and Rattner, 2002; Lucchesi et al., 2005). Recruitment of MSL complex to the X-chromosome is dependent on a recently discovered zinc finger protein called Chromatin-Linked Adapter for MSL Proteins (CLAMP) (Soruco et al., 2013).

We have previously demonstrated that CLAMP plays a key role in recruitment of the MSL complex to the X-chromosome in male *D. melanogaster* (Soruco et al., 2013). In addition to this role, we suggested that CLAMP has an essential function in general viability because targeting of *clamp* transcript by RNA interference results in a pupal lethal phenotype in both males and females. Further understanding of the *in vivo* function of CLAMP required a null mutant. However, due to the pericentric location of the *clamp* gene, no deficiencies or null mutations were available. We made several attempts at mutagenesis including P-element excision, homologous recombination, and sitedirected mutagenesis by Transcription Activator-Like Effector-based Nucleases (TALENs), which all proved to be unsuccessful. However, using the CRISPR/Cas9 system, we successfully introduced a nonsense mutation in the *clamp* gene that eliminates protein production. Our *clamp* null mutant revealed a dual role for CLAMP as an activator of dosage compensation in males and a repressor of important male-specific transcripts in females. Overall, we present a new tool for studying dosage compensation and suggest that CLAMP functions to assure that dosage compensation is sex-specific.

## Results and Discussion

### clamp^2^ mutants have a stronger lethal phenotype in males than in females

The *clamp* gene is located within pericentric heterochromatin on the left arm of chromosome two, one megabase from the centromere. Due to this chromosomal location, null mutants for the *clamp* gene were not previously available. Our attempts at generating a null mutation using well-established methods, including P-element excision, homologous recombination, and site-directed mutagenesis by TALENs all proved to be unsuccessful. We therefore used the CRISPR/Cas9 genome editing system, since one advantage of this approach is the potential introduction of nonsense mutations due to the resolution of double-stranded breaks by non-homologous end joining (Sander and Joung, 2014). We used the protein domain composition of CLAMP to determine where to target the Cas9 endonuclease. There are two predicted domains in CLAMP: an N-terminal glutamine-rich, low complexity domain, and a C-terminal zinc finger domain consisting of at least six zinc fingers (Figure 1A). Our laboratory previously demonstrated that the zinc finger domain of CLAMP is sufficient for DNA interactions (Soruco et al., 2013). Therefore, in order to generate a *clamp* null allele, we used the CRISPR/Cas9 system to target specifically upstream of the zinc finger domain of the *clamp* gene using the best available predicted guide RNAs.

**Figure 1:**
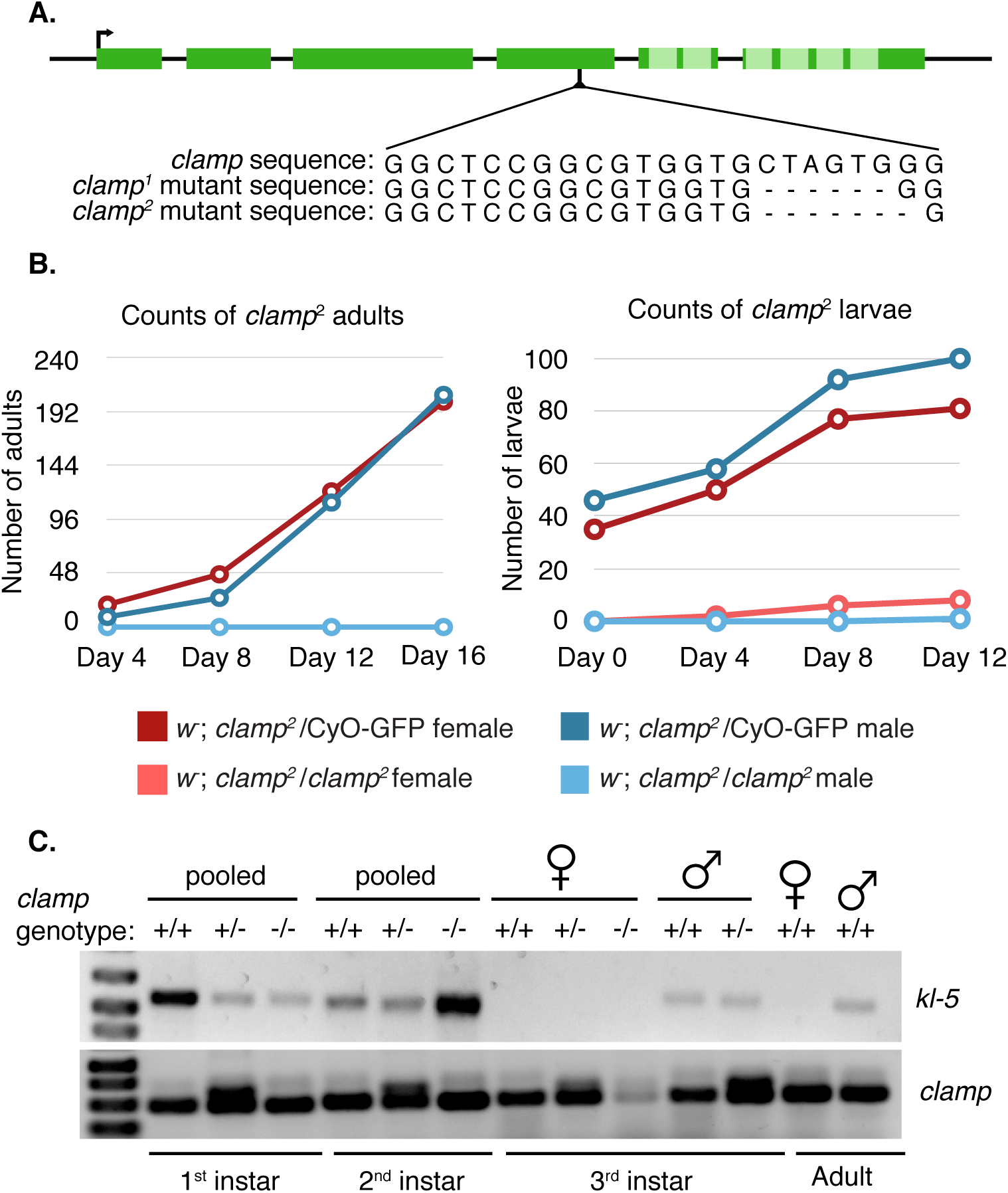
The *clamp*^*2*^ mutation have earlier lethality in males than in females. (A) The CRISPR/Cas9 introduced deletion is located in the fourth exon (dark green boxes) of the *clamp* gene, upstream of the DNA binding domain consisting of at least six zinc fingers (light green boxes). The homozygous viable *clamp*^*1*^ mutation is a six base pair deletion, while the homozygous lethal *clamp*^*2*^ mutation consists of the same six base pair deletion, with an additional seventh base removed. (B) The *clamp*^*2*^ mutation is homozygous lethal for both male and female adults (left). Heterozygous *clamp*^*2*^ male and female third instar larvae are viable, while homozygous *clamp*^*2*^ females are developmentally delayed. Homozygous *clamp*^*2*^ males are not viable. (C) The Y-chromosome gene *kl-5* is present in pooled homozygous *clamp*^*2*^ first and second instar larvae. Male third instar larvae are not viable, dying between the second and third instar larval stages.

Two different mutations were generated from this approach and balanced with a GFP-carrying CyO second chromosome balancer to track both larval and adult genotypes. Visual inspection of the wing phenotype in adult animals revealed that one mutation was homozygous viable while the other was not. Sequencing of the targeted region indicated that the homozygous viable animals carry a six base pair deletion in the *clamp* locus, resulting in the loss of two amino acids and an in-frame shift of the protein sequence (Figure 1A, *clamp*^*1*^). The homozygous lethal mutant carries the same six base pair deletion with an additional seventh base deleted (Figure 1A, *clamp*^*2*^). The seven base pair deletion causes a frameshift in the amino acid sequence, leading to an early stop codon occurring 13 amino acids after the mutation.

In order to quantify our observation that the *clamp*^*2*^ allele is adult homozygous lethal, we scored wing phenotype and counted the number of emerging homozygous (straight-winged) and heterozygous (curly-winged) *clamp*^*2*^ male and female adult flies. There is no apparent difference in the number of heterozygous *clamp*^*2*^ male or female animals (Figure 1B, left, red and dark blue lines). Consistent with a homozygous lethal mutation, we were unable to find any homozygous *clamp*^*2*^ mutants of either sex (Figure 1B, left panel). We next determined the viability of homozygous *clamp*^*2*^ larvae by scoring for the presence (in heterozygotes) or absence (in homozygotes) of GFP. We determined that homozygous *clamp*^*2*^ female larvae survive until third instar larval stages (Figure 1B, right panel, pink line). However, these larvae are developmentally delayed and occur at frequencies much lower than expected for Mendelian ratios (*χ*^2^ = 73.68, p¡0.001, Supplemental Table 3). In contrast, we only observed one homozygous male third instar larva out of a total 190 larvae.

We were interested in assaying when during development the majority of homozygous *clamp*^*2*^ male larvae die since we only observed a single third instar male in the balanced stock. This was done by testing for the presence of *clamp*^*2*^ homozygous first or second instar male larvae. It is difficult to phenotypically sex larvae prior to third instar. Therefore, we developed a PCR assay to establish the presence of male larvae by amplification of a Y-chromosome gene called *kl-5*. We extracted genomic DNA from pooled first or second instar larvae that were either homozygous or heterozygous for the *clamp*^*2*^ null allele, as determined by GFP fluorescence. We were unable to detect the *kl-5* gene in this assay in larval stages after the second instar, indicating that male larvae die between the second and third instar developmental stages (Figure 1C). Because *clamp* mRNA is heavily maternally deposited (Celniker et al., 2009; Graveley et al., 2011), it is possible that the maternal contribution of *clamp* transcript from heterozygous *clamp*^*2*^ mothers is sufficient for the homozygous males to survive well beyond zygotic genome activation.

### The clamp^2^ mutation is a null allele

The frame shift in the *clamp*^*2*^ allele results in a homozygous lethal phenotype, which suggests the mutation could be a recessive loss-of-function mutation. To characterize the mutation further, we analyzed the affect of the *clamp*^*2*^ allele on *clamp* RNA and protein abundance. We measured the production of *clamp* transcript by an established qRT-PCR assay (Soruco et al., 2013). We found that there is no statistical difference in the abundance of *clamp* transcript in both *clamp*^*2*^ heterozygous and homozygous female larvae when normalized to a wild type *y*^*‐*^*w*^*‐*^ female control (Figure 2A, yellow and orange bars). Interestingly, *clamp* transcript is dramatically increased in heterozygous mutant males when normalized to a wild type *y*^*‐*^*w*^*‐*^ male control (Figure 2A, grey bars). This observation raises the possibility that CLAMP may autoregulate its own expression. If autoregulation occurs, a reduction of functional CLAMP protein would cause an increase in *clamp* transcript to compensate for the loss of one functional allele. We previously demonstrated that CLAMP occupies its own promoter at high levels, supporting the hypothesis that the CLAMP protein regulates its own expression (Soruco et al., 2013). It is possible that this mechanism occurs in males and not in females due to the strong requirement for CLAMP in the essential process of recruiting MSL complex to the male X (Soruco et al., 2013).

**Figure 2:**
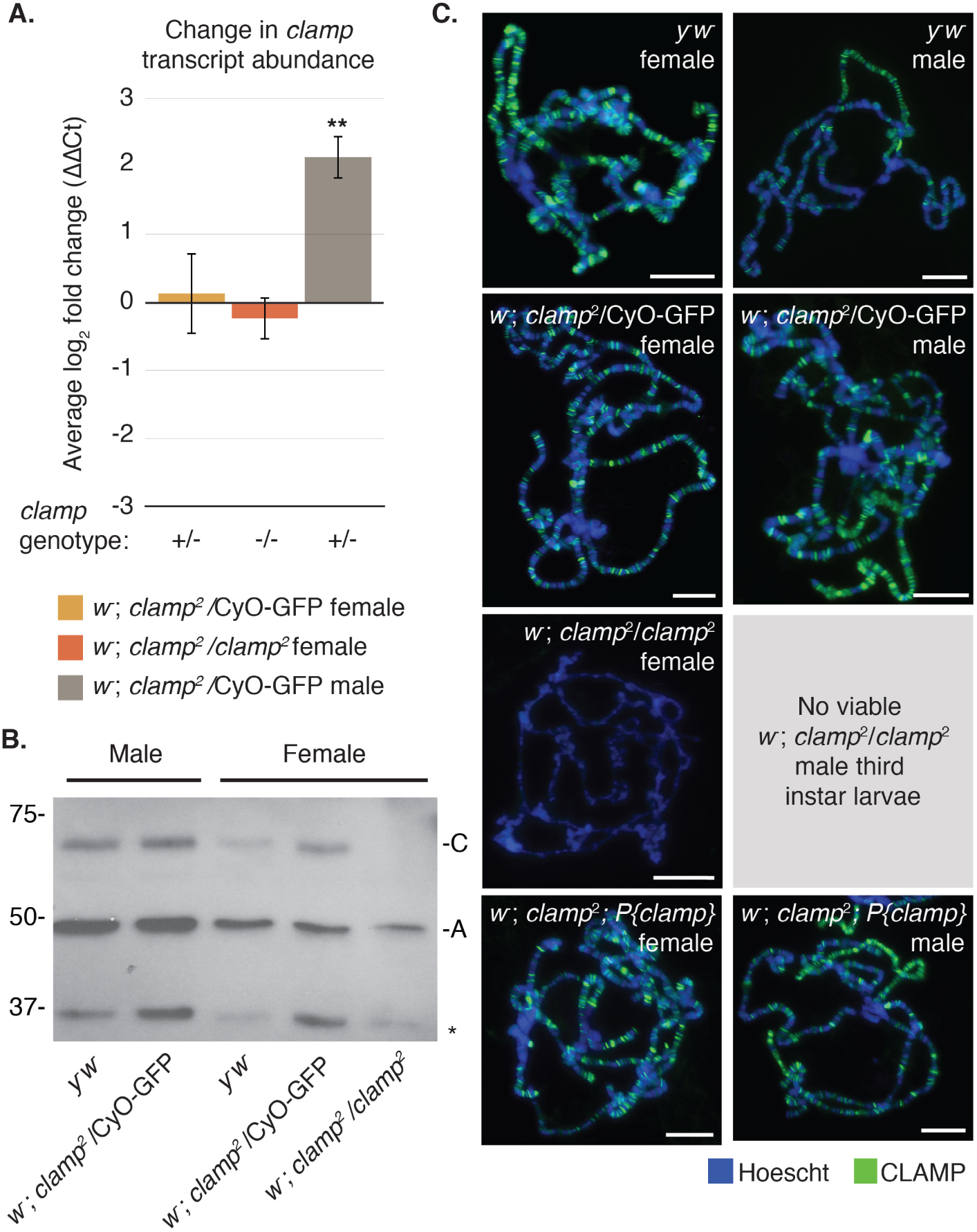
The *clamp*^*2*^ mutation is a null mutation.(A) Quantitative real-time PCR indicates no significant change in *clamp* transcript abundance in heterozygous or homozygous *clamp*^*2*^ female third instar larvae when normalized to *y*^*‐*^*w*^*‐*^ females. Males heterozygous for the *clamp*^*2*^ mutation show significant increase in *clamp* transcript compared to *y*^*‐*^*w*^*‐*^ males. Error bars are +/‐ 1 standard error of the mean (S.E.M., **p¡0.01, *p¡0.05). (B) Western blotting confirms CLAMP protein (C) is not produced in homozygous *clamp*^*2*^ females. Although there is a background band in all samples at 37kDA (*), a truncated form of CLAMP is not apparent as a result of the *clamp*^*2*^ mutation. Loading control is Actin (A). (C) Heterozygous *clamp*^*2*^ male and female larvae show no difference in CLAMP (green) localization on polytene chromosomes. There is no CLAMP localization in homozygous *clamp*^*2*^ female chromosomes. CLAMP immunostaining is rescued when a 12.5 Kb genomic region encompassing the *clamp* gene is inserted into the third chromosome. Scale bars are 0.02mm.

Although we did not observe a change in *clamp* mRNA levels in females, the frameshift deletion is located in the fourth exon of the gene, which could potentially abolish the zinc finger domain of the *clamp* protein. We thus hypothesized that we would observe an effect at the protein level in the form of a truncation. The CLAMP antibody epitope is located in the N-terminal region of the protein. Therefore, our antibody would detect a truncated form of CLAMP should one exist. To test this hypothesis, we performed western blot analysis using protein extracted from whole salivary glands of third instar larvae. The molecular weight of full length CLAMP is 61 kDa, whereas the *clamp*^*2*^ truncation is predicted to produce a 36.7 kDa protein. Western blotting indicates that although a background band is present in all samples around 37 kDa, a truncated form of CLAMP protein is not produced specifically in *clamp*^*2*^ mutants (Figure 2B). We have previously observed this background band and it is not ablated after *clamp* RNAi, suggesting that it is non-specific (Larschan et al., 2012). Moreover, western blotting reveals that homozygous *clamp*^*2*^ female larvae do not produce full-length CLAMP, despite producing wild type levels of *clamp* mRNA (Figure 2A).

To further test whether the *clamp*^*2*^ mutation is a null allele, we examined the localization of the CLAMP protein on polytene chromosomes. In wild type animals, CLAMP localizes throughout the genomes of both males and females (Figure 2C, *y*^*‐*^*w*^*‐*^ male and female). Furthermore, localization of CLAMP in heterozygous mutant males and females is not visibly distinct from that in wild type controls (Figure 2C, *w*^*‐*^; *clamp*^*2*^/CyO-GFP). In contrast, CLAMP staining in *clamp*^*2*^ homozygous female larvae indicated no localization of CLAMP to polytene chromosomes (Figure 2C, *w*^*‐*^; *clamp*^*2*^*/clamp*^*2*^). Homozygous male polytene chromosomes could not be obtained due to lethality of male third instar larvae (Figure 1B and C). Thus, polytene chromosome immunostaining indicates that CLAMP does not localize to chromatin in homozygous *clamp*^*2*^ females and further supports our conclusion that the *clamp*^*2*^ mutation is a null allele.

To determine if the homozygous lethality and loss of CLAMP localization is due to the frame shift mutation in the *clamp* gene rather than another background mutation, we generated a transgenic CLAMP rescue fly line. The stock contains a transgene that includes a 12.5kb region encompassing the *clamp* gene and all putative upstream regulatory regions but not any neighboring genes. The transgene is inserted into the VK33 attP docking site on the third chromosome (Venken et al., 2006). We found that *clamp*^*2*^ homozygous lethality is rescued in both male and female flies when one copy of the rescue construct is present. Moreover, the rescue construct generates a fully functional CLAMP protein that localizes to polytene chromosomes in both male and female homozygous *clamp*^*2*^ animals (Figure 2C, *w*^*‐*^; *clamp*^*2*^*; P{clamp*}).

### CLAMP represses roX RNA genes in females

CLAMP was initially identified as a transcription factor essential for directly linking the MSL complex to the X-chromosome in males (Larschan et al., 2012; Soruco et al., 2013). However, CLAMP also localizes throughout the genome and therefore has the potential to regulate additional transcripts (Figure 2C) (Soruco et al., 2013). Chromatin Immunoprecipitation followed by sequencing (ChIP-seq) demonstrates that CLAMP localizes to the 5 regulatory regions of genes that encode several MSL complex protein components (MSL1, MSL2, and MSL3) as well as the non-coding RNA components (*roX1* and *roX2*) (Soruco et al., 2013). We previously determined that CLAMP positively regulates the transcription of *roX2* in male (S2) *Drosophila* cells, likely because it recruits MSL complex to this genomic location (Soruco et al., 2013). However, the essential role of CLAMP in females remained unknown. Therefore, we compared the transcript abundance of MSL complex components in sexed wild type and *clamp*^*2*^ mutant third instar larvae.

We quantified transcript abundance of the following MSL complex component genes: *msl1*, *msl2*, *msl3*, *mle*, and *mof*. There was no significant difference in transcript abundance for any of the MSL complex components in heterozygous or homozygous females (Figure 3A, orange and yellow bars). However, we found significant activation of *msl1*, *msl3*, and *mof* in the *clamp*^*2*^ heterozygous mutant male animals compared to wild type *y*^*‐*^*w*^*‐*^ control males (Figure 3A, grey bars). We could not assay the levels of transcript in homozygous *clamp*^*2*^ mutant males because these males do not survive until third instar (Figure 1B and C). It is unclear whether CLAMP directly regulates transcription of MSL components in males or if the increase in MSL transcript levels in *clamp*^*2*^ heterozygotes is a result of a general role in transcription regulation that would also explain the increase in *clamp* transcript in these same animals (Figure 2A).

**Figure 3:**
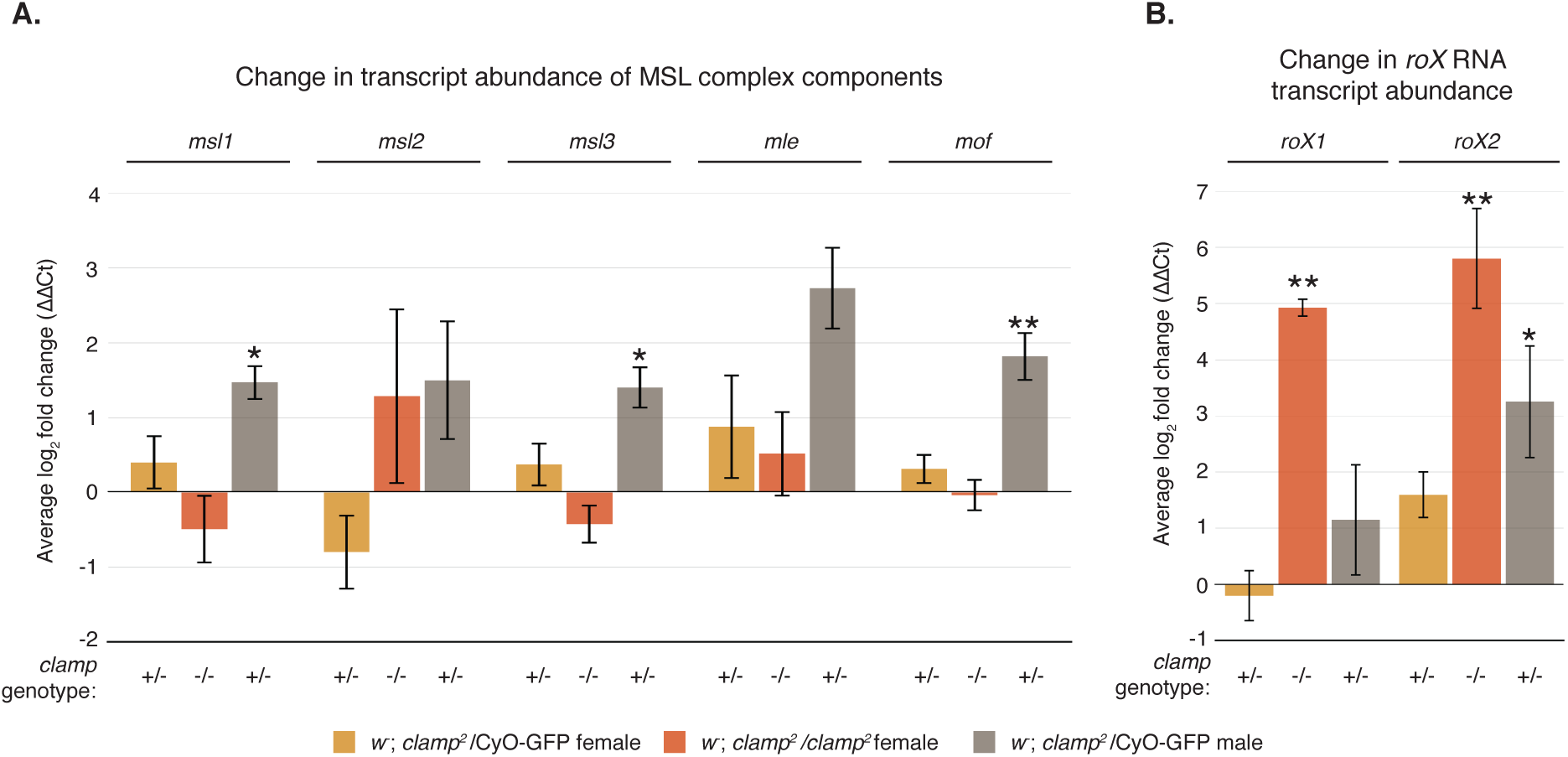
CLAMP regulates transcript abundance of MSL complex components. (A) Quantitative real-time PCR assay reveals CLAMP regulates transcripts of some MSL complex components in male larvae. There are significant increases in abundance for *msl1, msl3*, and *mof* in heterozygous males compared to *y*^*‐*^*w*^*‐*^ males. Though there are no significant differences in any MSL component transcript abundance in mutant females compared to *y*^*‐*^*w*^*‐*^ females, *msl2* transcript is highly variable in the homozygous *clamp*^*2*^ females. (B) CLAMP regulates expression of the *roX* RNAs in females and males as measured by qRT-PCR and normalized to respective *y*^*‐*^*w*^*‐*^ controls. The loss of CLAMP in homozygous *clamp*^*2*^ females results in a significant increase in the *roX* RNAs compared to *y*^*‐*^*w*^*‐*^ females. (Error bars are +/‐ 1 S.E.M., **p¡0.01, *p¡0.05).

Next, we tested whether loss of CLAMP affects production of the two non-coding RNA components of the MSL complex, *roX1* and *roX2*. In wild type female larvae, the *roX* RNAs are not expressed due to the absence of MSL complex, which activates their transcription (Meller et al., 1997; Meller, 2003). Interestingly, there is a dramatic and significant increase in the amount of both *roX1* and *roX2* transcript in homozygous *clamp*^*2*^ female larvae, which carry no functional copy of *clamp* (Figure 3B, orange bars), compared to *clamp*^*2*^ heterozygous females (Figure 3B, yellow bars). Similar to some MSL complex component transcripts, we observed an increase in *roX1* and *roX2* abundance in males heterozygous for the *clamp*^*2*^ mutation (Figure 3B, grey bars) when normalized to *y*^*-*^ *w*^*-*^ males. Therefore, CLAMP strongly represses the expression of male-specific *roX* RNA transcripts in females.

### The male form of Sxl transcript is present in clamp^2^ females

Formation and stability of the MSL complex is dependent on the presence of MSL2 protein (Kelley et al., 1995; Copps et al., 1998). Expression of MSL2 is regulated in a sex dependent way via de-stabilization of the *msl2* transcript by the master regulator protein Sex-lethal (Sxl), which links sex determination and MSL complex-dependent dosage compensation (Kelley et al., 1997; Bashaw and Baker, 1997). Production of Sxl protein is established early during development in XX females and maintained through autoregulatory mechanisms (Salz and Erickson, 2010). In females, Sxl inhibits translation of MSL2, preventing MSL complex formation. Sxl protein is not expressed in males, which stabilizes *msl2* transcripts to allow for MSL2 translation and subsequent MSL complex formation (Bashaw and Baker, 1997; Gebauer et al., 1998). Interestingly, transcript abundance of *msl2* is highly variable in homozygous *clamp*^*2*^ females and heterozygous *clamp*^*2*^ males (Figure 3A, orange and grey bars). This led us to hypothesize that the high variability is due to instability of *msl2* transcript, possibly as a result of misregulation of functional Sxl protein.

To test if CLAMP influences production of Sxl, we measured abundance of *Sxl* transcript. We observed no statistical difference in the amount of *Sxl* transcript present in either heterozygous and homozygous *clamp*^*2*^ females (Figure 4A, yellow and orange bars), when normalized to wild type *y*^*‐*^ *w*^*‐*^ females. Interestingly, there is a significant increase in the amount of *Sxl* transcript in *clamp*^*2*^ heterozygous males compared to wild type *y*^*‐*^*w*^*‐*^ males (Figure 4A, grey bars). It is well established that expression of Sxl protein is regulated through mechanisms of alternative splicing (Bell et al., 1988; Salz et al., 1989; Samuels et al., 1991). Thus, although the qRT-PCR results indicate no significant differences in the total amount of *Sxl* transcript in mutant female animals, this approach does not measure the relative abundance of specific splice forms.

**Figure 4:**
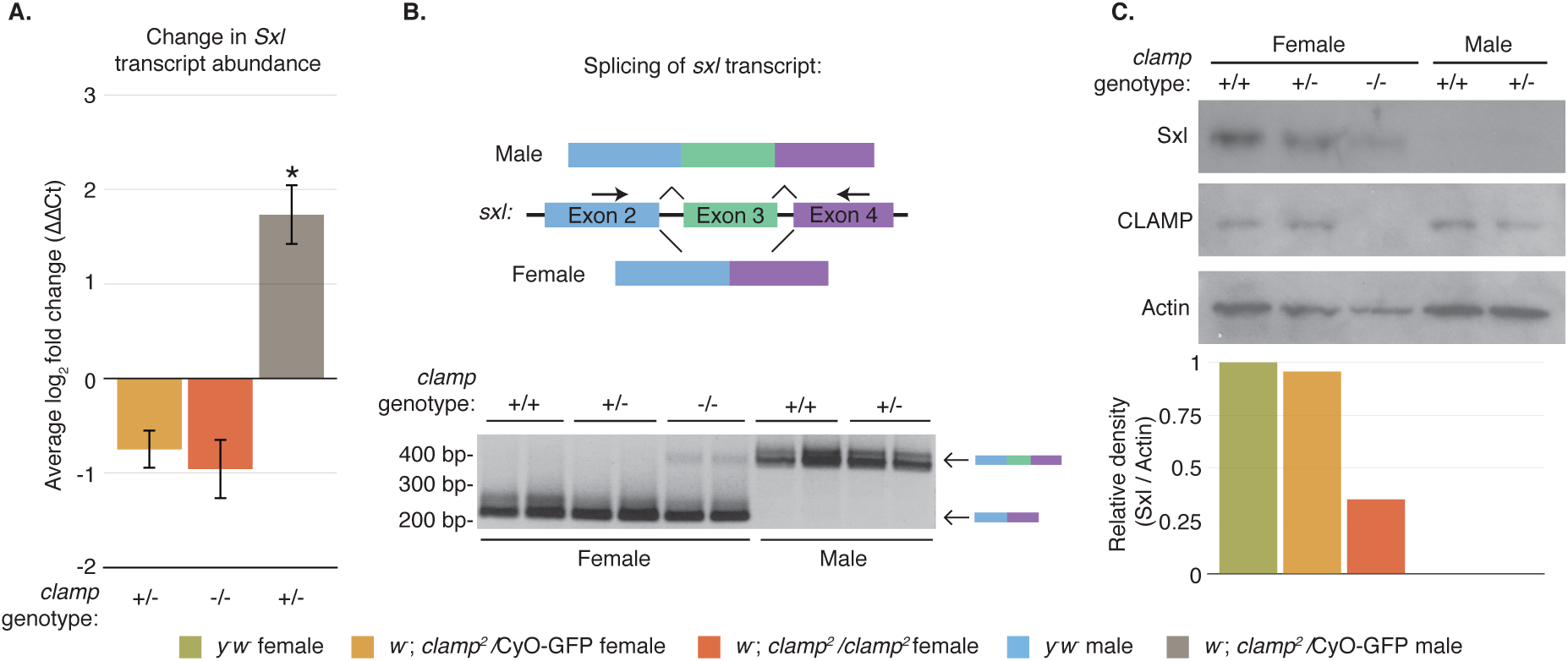
CLAMP regulates expression of Sex-lethal, the master regulator of sex determination. (A.) Expression of *sxl* transcript is significantly increased in heterozygous *clamp*^*2*^ males, when normalized to *y*^*‐*^*w*^*‐*^ males. (Error bars are +/‐ 1 S.E.M., **p¡0.01, *p¡0.05). (B) The female form of *sxl* transcript is alternatively spliced to exclude the third exon, resulting in a shorter PCR product than the male form of the transcript (top). Both male and female heterozygous *clamp*^*2*^ mutants correctly splice *sxl* transcript. However, the male form of the *sxl* transcript is aberrantly spliced in homozygous *clamp*^*2*^ females (bottom). For all genotypes, each lane is a separate replicate. (C). Presence of the non-functional male form of *sxl* transcript in homozygous *clamp*^*2*^ females is corroborated by a reduction in Sxl protein. The density of Sxl is normalized to Actin and plotted relative to *y*^*‐*^*w*^*‐*^ females.

*Sex-lethal* transcript is alternatively spliced to produce numerous spliceoforms. However, all female transcripts exclude the third exon, which contains a premature stop codon. *Sxl* transcript is produced in males, but the male-specific spliceoform includes the third exon to prevent functional protein from being produced (Bell et al., 1988; Salz et al., 1989; Samuels et al., 1991). Therefore, we designed a PCR assay to distinguish between the female (excluding exon 3) and male (including exon 3) versions of the *sxl* transcript (Figure 4B, left). Unexpectedly, we found that in homozygous *clamp*^*2*^ female animals, there is a small but detectable amount of the male (long) form of *Sxl* transcript (Figure 4B, top). These data reveal a novel role for CLAMP in the sex-determination pathway where CLAMP functions to prevent aberrant expression of the male specific *Sxl* transcript in females. Thus, although our qRT-PCR results indicate no significant reduction in the total amount of *Sxl* transcript in homozygous *clamp*^*2*^ females, the results of our splicing assay present the possibility that total functional transcript could be decreased as some transcripts contain exon 3.

To determine how the population of *Sxl* transcripts that include exon 3 in homozygous *clamp*^*2*^ females affects overall Sxl protein levels, we performed western blotting. We found a reduction in the amount of Sxl protein in the female homozygous *clamp*^*2*^ null mutants (Figure 4C, lane 3, orange bar) compared to wild type *y*^*‐*^*w*^*‐*^ and heterozygous females (lane 1, green bar and lane 2, yellow bar, respectively). We detected no Sxl protein in the male samples, as expected, because this protein is not expressed in males.

### Ectopic MSL complex does not form in clamp^2^ homozygous females

The *roX1* and *roX2* transcripts are not expressed in wild type females (Meller et al., 1997; Meller, 2003). Thus, the observed increase in abundance of these transcripts in *clamp*^*2*^ homozygous female larvae led us to question how this increase compared to wild type *roX* RNA expression in males. To test this, we reanalyzed *roX* RNA abundance (Figure 3B) by normalizing all transcript levels to *y*^*‐*^*w*^*‐*^ males. We found that while abundance of the *roX* RNAs are significantly depleted in *y*^*‐*^*w*^*‐*^ and heterozygous *clamp*^*2*^ females, abundance levels between *clamp*^*2*^ homozygous mutant females and *y*^*‐*^*w*^*‐*^ males are indistinguishable (Figure 5A, orange bars). Interestingly, as the number of functional copies of the *clamp* gene decreases, transcript abundance for *roX2* in females increases in a dose dependent manner. These data lead us to hypothesize that the homozygous *clamp*^*2*^ females could be dying due to an increase in *roX* expression and ectopic formation of MSL complex, which is known to be lethal in females (Kelley et al., 1995).

**Figure 5:**
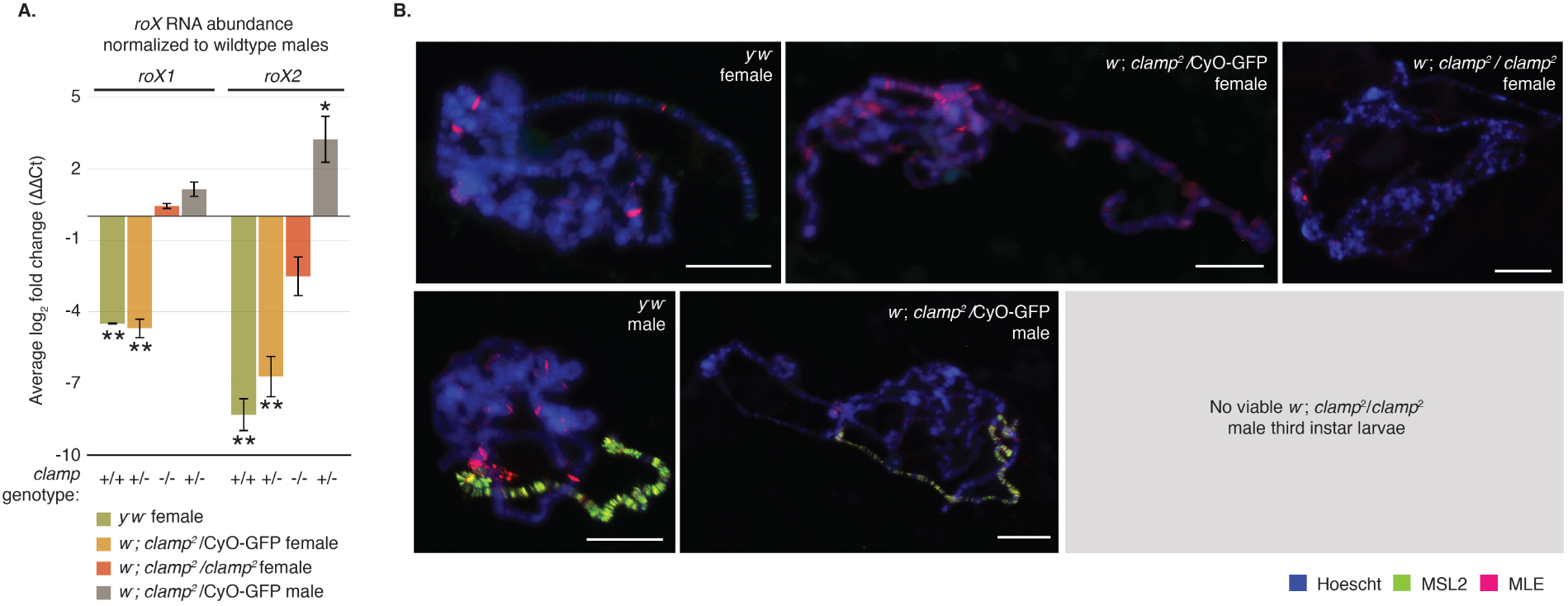
Ectopic MSL complex is not formed in *clamp*^*2*^ females. (A) *roX1* and *roX2* transcript abundance normalized to wild type males shows dramatic increase in expression in homozygous *clamp*^*2*^ females. (Error bars are +/‐ 1 S.E.M., **p¡0.01, *p¡0.05). (B) Polytene chromosome immunostaining shows that the core MSL complex component MSL2 is expressed only in males and localizes to the X-chromosome (green). The accessory protein, MLE, is expressed in both males and females and localizes throughout the genome (red). There is no change in localization of these proteins in heterozygous or homozygous *clamp*^*2*^ mutants. Scale bars are 0.02mm.

To test whether lethality in *clamp*^*2*^ homozygous females is caused by the increase in *roX* expression (Figure 5A), we generated a triple mutant fly line that is null for both *roX* genes in addition to *clamp. RoX* null females are usually viable. However, we found that the combined loss of both *roX* genes does not rescue the homozygous lethality of the *clamp*^*2*^ allele (data not shown). Thus we conclude that the increased expression of *roX* RNAs is not responsible for the lethality seen in *clamp*^*2*^ homozygous females.

It is known that MSL complex positively influences the expression of *roX* (Rattner and Meller, 2004). Moreover, ectopic expression of MSL complex is a molecular phenotype known to be lethal in females (Kelley et al., 1995). We observed reduced Sxl protein in *clamp*^*2*^ homozygous females, which could lead to stabilization of *msl2* transcript and subsequent MSL complex formation. Therefore, we asked whether lethality in the *clamp*^*2*^ homozygous females is due to ectopic formation of MSL complex, which would also explain the observed increased abundance of *roX* RNAs. We tested for ectopic formation of MSL complex in *clamp*^*2*^ homozygous females by immunostaining polytene chromosomes for localization of the MSL2 and MLE components of the MSL complex. We found there to be no evidence of ectopic localization of MSL2 in the heterozygous or homozygous mutant female animals (Figure 5B). However, this is not surprising, since we have previously demonstrated CLAMP is required for proper MSL complex recruitment (Larschan et al., 2012; Soruco et al., 2013). Therefore, if MSL complex forms in *clamp*^*2*^ homozygous females, it is likely that it would not be capable of localizing to the X-chromosome. Additionally, we see minimal differences in localization of MLE, a non-sex specific complex component known to be expressed in females and to have functions distinct from its role in dosage compensation (Figure 5B) (Cugusi et al., 2015).

Collectively, our results highlight the sexually dimorphic roles of the essential protein CLAMP. CLAMP was initially identified as a transcription factor necessary for recruiting MSL complex to the male X-chromosome. Here we demonstrate that CLAMP also functions to repress the expression of several key male-specific transcripts in females: the *roX* RNAs and the male-specific form of the *sxl* transcript. Importantly, we show that CLAMP regulates both the essential sex determination and dosage compensation pathways providing a new link between these two processes. The differential lethality of the *clamp*^*2*^ allele in males and females may be due to distinct roles for CLAMP in promoting dosage compensation in males and repressing male-specific transcripts in females.

In conclusion, CLAMP promotes sex determination in both males and females. Because CLAMP is heavily maternally deposited and enriched on the X-chromosome independent of MSL complex, it is possible that CLAMP is an important early regulator of sex determination. For example, it is possible that early in development of XX females, CLAMP influences proper splicing of *Sex-lethal* and represses *roX* RNA expression to ensure female fate, an exciting area for future investigation. In contrast, CLAMP performs the essential function of recruiting the MSL complex to the X-chromosome in males (Soruco et al., 2013). Overall, we demonstrate that CLAMP is a crucial regulator that promotes expression of male-specific transcripts in males and represses their expression in females.

## Materials and Methods

### Generation and validation of clamp mutant fly line

We used the FlyCRISPR Optimal Target Finder tool available from the University of Wisconsin to design a CRISPR target sequence for *clamp* (Gratz et al., 2014). We cloned target sequence oligonucleotides for *clamp* (sense: 5-CTT CGG CTC CGG CGT GGT GCT AGT-3 and antisense: 5-AAA CAC TAG CAC CAC GCC GGA GCC-3) into the pU6-BbsI-chiRNA plasmid (Addgene #45946) following the protocol outlined on the FlyCRISPR website. We validated correct ligation of the *clamp* CRISPR target sequence into the pU6-BbsI-chiRNA plasmid by Sanger sequencing using universal M13 primers.

The commercial service Genetic Services, Inc. microinjected the validated pU6-BbsI-chiRNA plasmid containing the *clamp* target sequence into germline-expressing Cas9 flies (*y*^*1*^, *w*^*1118*^;; PBac{vas-Cas9,U6-tracrRNA}VK00027). Flies containing a single mutation were returned balanced over the Curly of Oster (CyO) second chromosome balancer. We identified the CRISPR-generated mutation by PCR (Forward: 5-ACA ACT GAA GGG TTT GGA CGG-3, Reverse: 5-CAT GCA GGC TGA ACA AAC AG-3) followed by Sanger sequencing (Forward: 5-TCT GCA GGA CAA ACA CCT TG-3; Reverse: 5-CCC AAG CAC AAC TTC AGC AAA-3). We determined that the CRISPR/Cas9 system generated two *clamp* alleles: 1.) *y*^*1*^, *w*^*1118*^; *clamp*^*1*^ /CyO; and 2.) *y*^*1*^, *w*^*1118*^; *clamp*^*2*^*/CyO*.

### Generation and validation of clamp rescue fly line

We generated the CLAMP rescue construct using the P(acman) system that utilizes a conditionally amplifiable bacterial artificial chromosome (BAC) clone, recombineering, and bacteriophage phiC31 mediated insertion at a genomic attB site (Venken et al., 2006). We designed primers for two homology arms to capture a 12.5kb region spanning the entire clamp locus (3.5kb) including the presumed promoter (Left Homology Arm (1.2kb) Forward: 5-ACC GGC GCG CCG CAG AAG GAA GAG TTT CCG A-3, Reverse: 5‐ CGC GGA TCC AAG TCC TGG CCT AAG CCC TA-3; Right Homology Arm (800bp) Forward: 5‐ CGC GGA TCC TTT TGT GCA TGG TCA ACC ACG-3, Reverse: 5‐ ACC TTA ATT AAG GGC AAA CAT ATT TCG CAC GAT AC-3). We amplified homology arms off a conditionally amplifiable P(acman) BAC clone, Ch322 20C06 (BacPac Resources) using Copy Control (Epicenter) reagent for vector amplification. We simultaneously cloned the arms into the PacMan vector 3XP3-eGFP-attB-Amp (gift from Koen Venken) at the multicloning site (MCS) using the engineered restriction sites AscI-BamHI (left) and BamHI-PacI (right) in a three-component ligation. We identified positive colonies via Sanger sequencing across the MCS. Using BamH1, we linearized the intermediate vector and purified the product. Next, the linearized vector was transformed into *E.coli* that we had previously transformed with the *clamp* containing BAC clone Ch322 20C06 and expressing the mini-lambda vector encoding the phiC31 recombinase (SW102, NCI BRB Preclinical Repository). We identified positive colonies via sequencing across the left and right homology arm junction.

Genetic Services, Inc. microinjected the full rescue construct into *D. melanogaster* embryos containing the attB docking site (VK33) on Chromosome 3L band 65B2 (Venken et al., 2006). We identified the rescue construct line using eGFP expression, and maintained the subsequent stock in the homozygous state (*y*^*1*^, *w*^*1118*^;; P{3xP3-eGFP, *clamp* = *clamp*}).

### Fly crosses and counts

To generate a *clamp*^*2*^ mutant line with a larval phenotypic marker, we used standard methods to cross the original *y*^*1*^, *w*^*1118*^; *clamp*^*2*^/CyO stock to a CyO-GFP stock that expresses GFP at all stages of larval development (*w*^*1118*^; *sna*^*Sco*^/CyO, P{ActGFP.w^-^}CC2). The resulting *w*^*1118*^; *clamp*^*2*^/CyO-GFP stock (referred to as *clamp*^*2*^ in text) contains both larval and adult phenotypic markers and was used for all remaining experiments. To assess larval viability, we collected 190 third instar larvae over the course of 12 days. For each larva, we determined the sex and *clamp* genotype. From these larvae, we monitored eclosion of the pupae into adult flies every day and totaled every four days for 16 days.

To test if the *clamp*^*2*^ mutation can be rescued, we crossed the *w*^*1118*^; *clamp*^*2*^/CyO-GFP stock to the *y*^*1*^, *w*^*1118*^;; P{3xP3-eGFP, *clamp*} rescue line and scored viability at the adult stage by wing phenotype.

### Sample collection and PCR for kl-5 gene

We tested for the presence of the Y-chromosome gene *kl-5* in first, second, and third instar larvae of the following animals: 1. *y*^*1*^, *w*^*1118*^, 2. *w*^*1118*^; *clamp*^*2*^/CyO-GFP, and 3. *w*^*1118*^; *clamp*^*2*^/*clamp*^*2*^. We collected and pooled 10 larvae each of the first and second instar developmental stage. For third instar larvae, we dissected 10 salivary glands of sexed males and females of each genotype in cold PBS. As an additional control, we tested 10 adult male and adult female whole flies. We flash froze all samples in liquid nitrogen and homogenized using a bead mill. Next, we suspended the homogenized samples in 30uL of lysis buffer (10mM Tris-HCl pH8.0, 1mM EDTA, 25mM NaCl, 0.2mg/ml Proteinase K, 1ng/uL RNase) and incubated at 37C for 30 minutes, followed by a 5 minute incubation at 90C. We purified genomic DNA by standard phenol:chloroform extraction using Phase-lock tubes (5 Prime) as per the manufacturers instructions followed by ethanol precipitation.

We tested purified genomic DNA for the presence of the *kl-5* gene by PCR using the following primers (Forward: 5-ATC GCA AAC GAG TGG TCT CA-3; Reverse: 5-TGT ATC AAG GGC AGG CAT CC-3). As a genomic DNA loading control, we amplified the *clamp* locus with the PCR primers used to identify the mutation.

### Quantitative Real Time PCR (qRT-PCR)

To analyze transcript abundance, we used TRIzol (Thermo Fisher Scientific) per the manufacturers instructions to extract total RNA from three biological replicates of five third instar larvae from each genotype. We reverse-transcribed one microgram of total RNA using the SuperScript VILO cDNA Synthesis Kit (Life technologies) by following the manufacturers protocol. Three technical replicates for each target transcript were amplified using SYBR Green (Life Technologies) on an Applied Biosystems StepOnePlus™ Real-Time PCR System. Primers were used at a concentration of 200nM to amplify targets from 2ng of cDNA. Primer sequences for qRT-PCR are in Supplemental Table 1. We calculated transcript abundance using the standard ΔΔ Ct method with GAPDH as the internal control (Livak and Schmittgen, 2001). We normalized female mutant samples to the female *y*^*1*^, *w*^1118^(referred to as *y*^*‐*^ *w*^*‐*^ in text) control, except where specified. We normalized male mutant samples to the male *y*^*1*^, *w*^*1118*^ control. We tested statistical significance by performing an ANOVA multiple comparison test on the mean ΔCt values, followed by a Tukey post hoc analysis for multiple comparison correction. F-statistics and p-values for all comparisons are presented in Supplemental Table S2.

To test for the male and female forms of the *Sex-lethal* transcript, we extracted total RNA from two biological replicates of ten third instar larvae from each genotype. We reverse-transcribed two micrograms of total RNA using the SuperScript VILO cDNA Synthesis Kit (Life technologies) following the manufacturers protocol. We amplified *sex-lethal* transcript by PCR using primers designed to span the exon 2 exon 4 junction (Exon 2 Forward: 5-TGC AAC TCA CCT CAT CAT CC-3; Exon 4 Reverse: 5-GAT GGC AGA GAA TGG GAC AT-3) (Figure 4B).

### Western blotting

We dissected salivary glands from third instar larvae in cold PBS and froze samples in liquid nitrogen. We extracted total protein from the samples by homogenizing in lysis buffer (50mM Tris-HCl pH 8.0, 150mM NaCl, 1% SDS, 0.5X protease inhibitor) using a small pestle. After a five-minute incubation at room temperature, we cleared the samples by centrifuging at room temperature for 10 minutes at 14,000 x g. To blot for CLAMP and Actin, we ran 5 micrograms of total protein on a Novex 10% Tris-Glycine precast gel (Life technologies). To measure Sex-lethal protein levels, we ran 20 micrograms total protein on a Novex 12% Tris-Glycine precast gel (Life technologies). We transferred proteins to PVDF membranes using the iBlot transfer system (ThermoFisher Scientific) and probed the membranes for CLAMP (1:1000, SDIX), Actin (1:400,000, Millipore), and Sex-lethal (1:500, gift from Fatima Gebauer) using the Western Breeze kit following the manufacturers protocol (ThermoFisher Scientific).

Relative expression of Sex-lethal protein was quantified using the gel analysis tool in ImageJ software following the guidelines outlined on the website (Schneider et al., 2012). For each genotype, we first internally normalized the amount of Sex-lethal protein to Actin. Next, we determined relative expression of Sex-lethal by comparing the Actin normalized quantities to *y*^*1*^, *w*^*1118*^ female samples.

### Polytene chromosome squashes and immunostaining

We prepared polytene chromosome squashes as previously described (Cai et al., 2010). We stained polytene chromosomes with anti-CLAMP (1:1000, SDIX), anti-MLE (1:500), or anti-MSL2 (1:500) primary antibodies. For detection, we used all Alexafluor secondary antibodies at a concentration of 1:1000. We visualized slides at 40X on a Zeiss Axioimager M1 Epifluorescence upright microscope with the AxioVision version 4.8.2 software.

## Data Availability

Generated fly stocks are available upon request. Supplemental Table 1 contains primer sequences for qRT-PCR assays.

## Acknowledgements

Thank you to Dr. Fatima Gebauer Hernandez from the Centre for Genomic Regulation in Barcelona, Spain for gifting us a generous portion of the anti-Sex-lethal antibody. Also, thank you to Dr. Koen Venkin for providing us with the PacMan vector. We would also like to thank John Urban for valuable input on experimental analysis and critical review of the manuscript. J.A.U. was supported by National Institutes of Health award F31GM108423. All experiments were supported by National Institutes of Health R01GM098461-1, American Cancer Society Research Scholar 123682-RSG-13-040-01-DMC, and a Pew Biomedical Scholars program grant.

## References

Bashaw, G. J. and B. S. Baker 1997. The regulation of the Drosophila msl-2 gene reveals a function for Sex-lethal in translational control. Cell, 89(5):789–98.

Bell, L. R., E. M. Maine, P. Schedl, and T. W. Cline 1988. Sex-lethal, a Drosophila sex determination switch gene, exhibits sex-specific RNA splicing and sequence similarity to RNA binding proteins. Cell, 55(6):1037–1046.

Cai, W., Y. Jin, J. Girton, J. Johansen, and K. M. Johansen 2010. Preparation of Drosophila polytene chromosome squashes for antibody labeling. Journal of visualized experiments: Jo VE, (36).

Celniker, S. E., L. A. L. Dillon, M. B. Gerstein, K. C. Gunsalus, S. Henikoff, G. H. Karpen, M. Kellis, E. C. Lai, J. D. Lieb, D. M. MacAlpine, G. Micklem, F. Piano, M. Snyder, L. Stein, K. P. White, and R. H. Waterston 2009. Unlocking the secrets of the genome. Nature, 459(7249):927–30.

Copps, K., R. Richman, L. M. Lyman, K. A. Chang, J. Rampersad-Ammons, and M. I. Kuroda 1998. Complex formation by the Drosophila MSL proteins: role of the MSL2 RING finger in protein complex assembly. The EMBO journal, 17(18):5409–17.

Cugusi, S., S. Kallappagoudar, H. Ling, and J. C. Lucchesi 2015. The Drosophila Helicase Maleless (MLE) is Implicated in Functions Distinct From its Role in Dosage Compensation. Molecular & cellular proteomics: MCP, 14(6):1478–88.

Gebauer, F., L. Merendino, M. W. Hentze, and J. Valcárcel 1998. The Drosophila splicing regulator sex-lethal directly inhibits translation of male-specific-lethal 2 mRNA. RNA (New York, N.Y.), 4(2):142–50.

Gratz, S. J., F. P. Ukken, C. D. Rubinstein, G. Thiede, L. K. Donohue, A. M. Cummings, and K. M. O’Connor-Giles 2014. Highly specific and efficient CRISPR/Cas9-catalyzed homology-directed repair in Drosophila. Genetics, 196(4):961–71.

Graveley, B. R., A. N. Brooks, J. W. Carlson, M. O. Duff, J. M. Landolin, L. Yang, C. G. Artieri, M. J. van Baren, N. Boley, B. W. Booth, J. B. Brown, L. Cherbas, C. A. Davis, A. Dobin, R. Li, W. Lin, J. H. Malone, N. R. Mattiuzzo, D. Miller, D. Sturgill, B. B. Tuch, C. Zaleski, D. Zhang, M. Blanchette, S. Dudoit, B. Eads, R. E. Green, A. Hammonds, L. Jiang, P. Kapranov, L. Langton, N. Perrimon, J. E. Sandler, K. H. Wan, A. Willingham, Y. Zhang, Y. Zou, J. Andrews, P. J. Bickel, S. E. Brenner, M. R. Brent, P. Cherbas, T. R. Gingeras, R. A. Hoskins, T. C. Kaufman, B. Oliver, and S. E. Celniker 2011. The developmental transcriptome of Drosophila melanogaster. Nature, 471(7339):473–9.

Kelley, R. L., I. Solovyeva, L. M. Lyman, R. Richman, V. Solovyev, and M. I. Kuroda 1995. Expression of msl-2 causes assembly of dosage compensation regulators on the X chromosomes and female lethality in Drosophila. Cell, 81(6):867–77.

Kelley, R. L., J. Wang, L. Bell, and M. I. Kuroda 1997. Sex lethal controls dosage compensation in Drosophila by a non-splicing mechanism. Nature, 387(6629):195–9.

Larschan, E., M. M. L. Soruco, O.-K. Lee, S. Peng, E. Bishop, J. Chery, K. Goebel, J. Feng, P. J. Park, and M. I. Kuroda 2012. Identification of chromatin-associated regulators of MSL complex targeting in Drosophila dosage compensation. PLoS genetics, 8(7):e1002830.

Livak, K. J. and T. D. Schmittgen 2001. Analysis of relative gene expression data using real-time quantitative PCR and the 2(-Delta Delta C(T)) Method. Methods (San Diego, Calif.), 25(4):402–8.

Lucchesi, J. C., W. G. Kelly, and B. Panning 2005. Chromatin remodeling in dosage compensation. Annual review of genetics, 39:615–51.

Meller, V. H. 2003. Initiation of dosage compensation in Drosophila embryos depends on expression of the roX RNAs. Mechanisms of Development, 120(7):759–767.

Meller, V. H. and B. P. Rattner 2002. The roX genes encode redundant male-specific lethal transcripts required for targeting of the MSL complex. The EMBO journal, 21(5):1084–91.

Meller, V. H., K. H. Wu, G. Roman, M. I. Kuroda, and R. L. Davis 1997. roX1 RNA Paints the X Chromosome of Male Drosophila and Is Regulated by the Dosage Compensation System. Cell, 88(4):445–457.

Rattner, B. P. and V. H. Meller 2004. Drosophila male-specific lethal 2 protein controls sex-specific expression of the roX genes. Genetics, 166(4):1825–32.

Salz, H. K. and J. W. Erickson 2010. Sex determination in Drosophila: The view from the top. Fly, 4(1):60–70.

Salz, H. K., E. M. Maine, L. N. Keyes, M. E. Samuels, T. W. Cline, and P. Schedl 1989. The Drosophila female-specific sex-determination gene, Sex-lethal, has stage-, tissue-, and sex-specific RNAs suggesting multiple modes of regulation. Genes & development, 3(5):708–19.

Samuels, M. E., P. Schedl, and T. W. Cline 1991. The complex set of late transcripts from the Drosophila sex determination gene sex-lethal encodes multiple related polypeptides. Molecular and cellular biology, 11(7):3584–602.

Sander, J. D. and J. K. Joung 2014. CRISPR-Cas systems for editing, regulating and targeting genomes. Nature biotechnology, 32(4):347–55.

Schneider, C. A., W. S. Rasband, and K. W. Eliceiri 2012. NIH Image to ImageJ: 25 years of image analysis. Nature Methods, 9(7):671–675.

Soruco, M. M. L., J. Chery, E. P. Bishop, T. Siggers, M. Y. Tolstorukov, A. R. Leydon, A. U. Sugden, K. Goebel, J. Feng, P. Xia, A. Vedenko, M. L. Bulyk, P. J. Park, and E. Larschan 2013. The CLAMP protein links the MSL complex to the X chromosome during Drosophila dosage compensation. Genes & development, 27(14):1551–6.

Venken, K. J. T., Y. He, R. A. Hoskins, and H. J. Bellen 2006. P[acman]: a BAC transgenic platform for targeted insertion of large DNA fragments in D. melanogaster. Science (New York, N.Y.), 314(5806):1747–51.

